# X-ray structures of human ET_B_ receptor provide mechanistic insight into receptor activation and partial activation

**DOI:** 10.1101/343756

**Authors:** Wataru Shihoya, Tamaki Izume, Asuka Inoue, Keitaro Yamashita, Francois Marie Ngako Kadji, Kunio Hirata, Junken Aoki, Tomohiro Nishizawa, Osamu Nureki

## Abstract

Endothelin receptors (ET_A_ and ET_B_) are class A GPCRs activated by vasoactive peptide endothelins, and are involved in blood pressure regulation. ET_B_-selective signaling induces vasorelaxation, and thus selective ET_B_ agonists are expected to be utilized for improved anti-tumour drug delivery and neuroprotection. The effectiveness of a highly ET_B_-selective endothelin analogue, IRL1620, has been investigated in clinical trials. Here, we report the crystal structures of human ET_B_ receptor in complex with ET_B_-selective agonists, endohelin-3 and IRL1620. The 2.0 Å-resolution structure of the endothelin-3-bound receptor revealed that the disruption of water-mediated interactions between W6.48 and D2.50, which are highly conserved among class A GPCRs, is critical for receptor activation. These hydrogen-bonding interactions are partially preserved in the IRL1620-bound structure, and a functional analysis revealed the partial agonistic effect of IRL1620. The current findings clarify the detailed molecular mechanism for the coupling between the orthosteric pocket and the G-protein binding, and the partial agonistic effect of IRL1620, thus paving the way for the design of improved agonistic drugs targeting ET_B_.

## Introduction

Endothelin receptors belong to the class A GPCRs, and are activated by endothelins, which are 21-amino acid peptide agonists^1^. Both of the endothelin receptors (the ET_A_ and ET_B_ receptors) are widely expressed in the human body, including the vascular endothelium, brain, lung, kidney, and other circulatory organs^2,3^. Three kinds of endothelins (ET-1, ET-2, and ET-3) activate the endothelin receptors (ETRs) with sub-nanomolar affinities. ET-1 and ET-2 show similar affinities to both of the endothelin receptors, while ET-3 shows two orders of magnitude lower affinity to ET_A_^4-6^. The stimulation of the ET_A_ receptor by ET-1 leads to potent and long-lasting vasoconstriction, whereas that of the ET_B_ receptor induces nitric oxide-mediated vasorelaxation^7-9^. The human brain contains the highest density of endothelin receptors, with the ET_B_ receptor comprising about 90% in areas such as the cerebral cortex^10^. The ET_B_ receptor in neurons and astrocytes has been implicated in the promotion of neuroprotection, including neuronal survival and reduced apoptosis^11,12^. Moreover, the ET-3/ET_B_ signaling pathway has distinct physiological roles, as compared to the ET-1 pathway. In the brain, ET-3 is responsible for salt homeostasis, by enhancing the sensitivity of the brain sodium-level sensor Nax channel^13^. The ET-3/ET_B_ signaling pathway is also related to the development of neural crest cells, and plays an essential role in the formation of the enteric nervous system^14^. Thus, mutations of the ET-3 or ET_B_ genes cause Hirschsprung’s disease, a birth defect in which nerves are missing from parts of the intestine^15,16^. Overall, the endothelin system participates in a wide range of physiological functions in the human body.

Since the activation of the ET_B_ receptor has a vasodilating effect, unlike the ET_A_ receptor, ET_B_-selective agonists have been studied as vasodilator drugs for the improvement of tumour drug delivery, as well as for the treatment of hypertension^2,3^. IRL1620 (N-Suc-[E9, A11, 15] ET-1_8-21_)^17^, a truncated peptide analogue of ET-1, is the smallest agonist that can selectively stimulate the ET_B_ receptor, and currently no non-peptidic ET_B_-selective agonists have been developed. The affinity of IRL1620 to the ET_B_ receptor is comparable to that of ET-1, whereas it essentially does not activate the ET_A_ receptor, and thus it shows high ET_B_ selectivity of over 100,000-fold. Due to its large molecular weight, IRL1620 is not orally active and thus requires intravenous delivery. Despite its pharmacokinetic disadvantages, IRL1620 is an attractive candidate for the treatment of various diseases related to the ET_B_ receptor. Since the ET_B_-selective signal improves blood flow, IRL1620 could be utilized for the improved efficacy of anti-cancer drugs by increasing the efficiency of drug delivery, as shown in rat models of prostate and breast cancer^18-21^. Moreover, this strategy can also be applied to radiotherapy in the treatment of solid tumours, as the raRef18diation-induced reduction in the tumour volume was enhanced by IRL1620^22^. IRL1620 also has vasodilation and neuroprotection effects in the brain. IRL1620 reduced neurological damage following permanent middle cerebral artery occlusion in a rat model of focal ischaemic stroke^23^. Moreover, the stimulation of the ET_B_ receptors by IRL1620 reduces the cognitive impairment induced by beta amyloid (1-40), a pathological hallmark of Alzheimer’s disease, in rat experiments^24,25^. These data suggest that ET_B_ selective agonists might offer new therapeutic strategies for neuroprotection and Alzheimer’s disease. The safety and maximal dose of IRL1620 were investigated in a phase I study. While a recent phase 2 study of IRL1620 in combination with docetaxel as the second-line drug reported no significant improvement in the treatment of advanced biliary tract cancer (ABTC)^26^, further trials for selected patients based on tumour types with various choices for the second-line drugs are still expected. Concurrently, the pharmacological properties of IRL1620 could be also improved for better clinical applications. However, little is known about the selectivity and activation mechanism of this artificially designed agonist peptide, although the ET_B_ structures in complex with ET-1 and antagonists have been determined^27,28^.

In this study, we report the crystal structures of the ET_B_ receptor in complex with two ET_B_-selective ET variant agonists, ET-3 and IRL1620, together with their detailed biochemical characterization. These results reveal the different activation mechanisms of these agonists, especially for the partial activation by IRL1620.

## Results

### Functional characterization of ET-3 and IRL1620

We first investigated the biochemical activities of ET-3 and IRL1620 for the human endothelin receptors, by TGFα shedding (G-protein activation) and β-arrestin recruitment assays. The EC_50_ and E_max_ values of ET-3 for the ET_B_ receptor were similar to those of ET-1 in both assays, while the EC_50_ value for ET_A_ was about 5-fold lower (Fig. 1a-d). These data indicate that ET-3 functions as a full agonist for the endothelin receptors, with moderate ET_B_-selectivity. The EC_50_ value of IRL1620 for the ET_B_ receptor was 0.11 nM in the TGFα shedding assay, and was almost same as that of ET-1 (Fig. 1b). By contrast, a 320 nM concentration of IRL1620 did not activate ET_A_ in the TGFα shedding assay (Fig. 1a). These data showed that IRL1620 is ET_B_-selective by over 3,000-fold, in excellent agreement with previous functional analyses^17,29^. However, despite its sub-nanomolar affinity, the E_max_ values of IRL1620 for the ET_B_ receptor were 88% (TGFα shedding assay) and 87% (β-arrestin recruitment assay) of those of ET-1 (Fig. 1b, d), suggesting that IRL1620 functions as a partial agonist for the ET_B_ receptor.

**Fig. 1.**
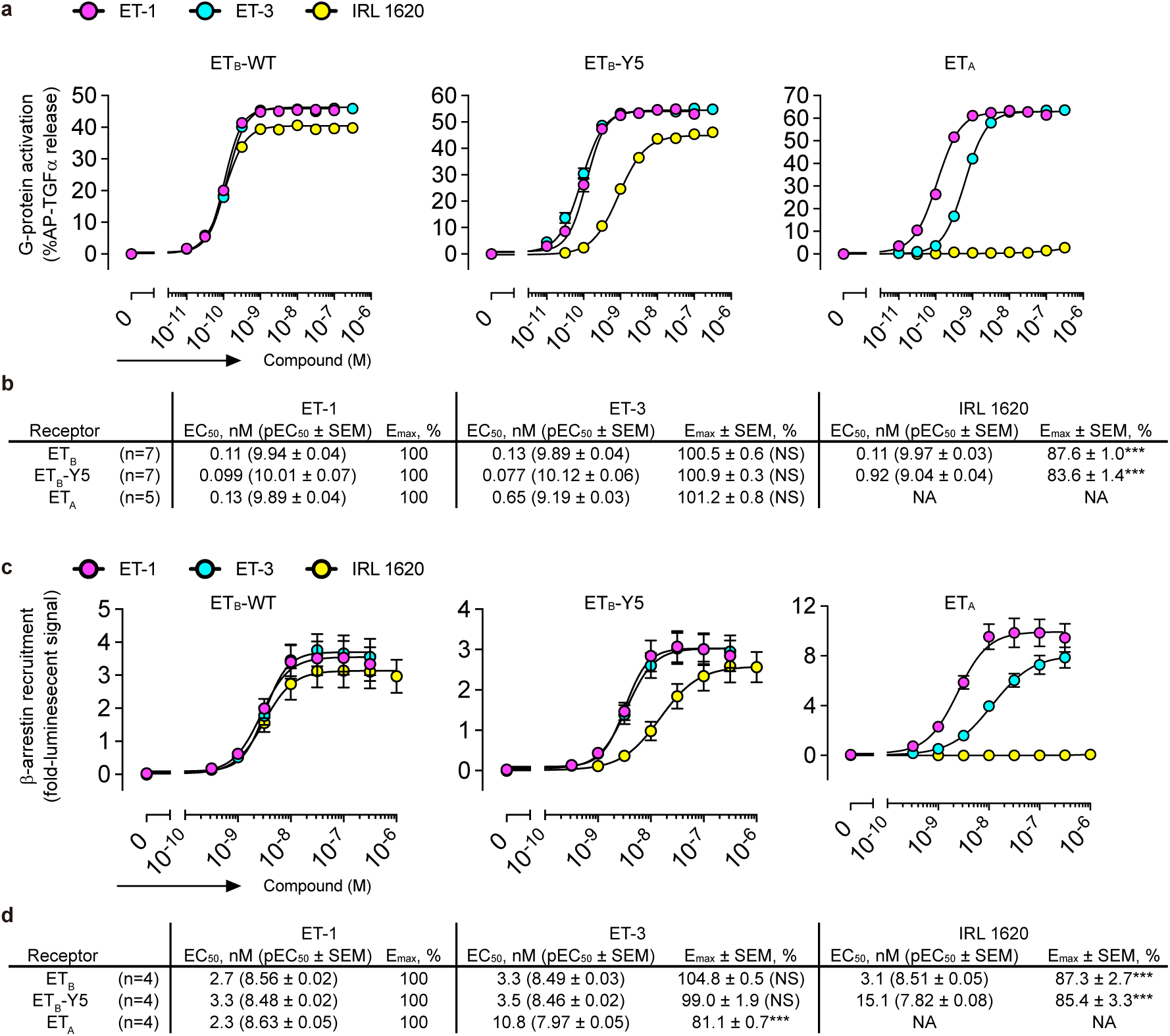
Pharmacological characterizations of ET-3 and IRL 1620. **a**, Concentration response-curves of AP-TGFα release in the ET-1, ET-3, and IRL1620 treatments of HEK293 cells expressing the endothelin receptors. Symbols and error bars are means and s.e.m. (standard error of the mean) of five or seven independent experiments, each performed in triplicate. Note that the error bars are smaller than the symbols for most data points. **b**, The EC_50_ and E_max_ values of the AP-TGFα release response for the endothelin receptors. **c**, Concentration response-curves of β-arrestin recruitment in the ET-1, ET-3, and IRL1620 treatments of HEK293 cells expressing the endothelin receptors. Symbols and error bars are means and s.e.m. of five independent experiments, each performed in duplicate. **d**, The EC_50_ and E_max_ values of the β-arrestin recruitment for the endothelin receptors. The E_max_ value of the β-arrestin recruitment in the ET-1 treatment was normalized to 100% for each experiment. E_max_ values that significantly differ from the wild-type are denoted by asterisks. \**P* < 0.05; ** *P* < 0.01; *** *P* < 0.001, one-way ANOVA with Dunnett’s post hoc test. NS, not significant.

To obtain mechanistic insights into the different actions of these agonists, we performed X-ray crystal structural analyses of the human ET_B_ receptor in complex with ET-3 and IRL1620. For crystallization, we used the previously established, thermostabilized ET_B_ receptor (ET_B_-Y5)^27,30^. We confirmed that the thermostabilizing mutations minimally affect the binding of these agonists. The EC_50_ and E_max_ values of ET-3 and IRL1620 for ET_B_-Y5 were comparable to those for the wild type receptor in both assays (Fig. 1b, d). In addition, IRL1620 also functions as a partial agonist for the thermostabilized receptor, as the E_max_ values for ET_B_-Y5 were lower than those of ET-1 in both assays (84% and 85% in the TGFα shedding assay and the β-arrestin recruitment assay, respectively). To facilitate crystallization, we replaced the third intracellular loop (ICL3) of the receptor with T4 Lysozyme (ET_B_-Y5-T4L), and using *in meso* crystallization, we obtained crystals of ET_B_-Y5-T4L in complex with ET-3 and IRL1620 (Supplementary Fig. 1a, b). In total, 757 and 68 datasets were collected for the ET-3- and IRL1620-bound receptors, respectively, and merged by a data processing system KAMO^31^. Eventually, we determined the ET_B_ structures in complex with ET-3 and IRL1620 at 2.0 and 2.7 Å resolutions, respectively, by molecular replacement using the ET-1-bound receptor (PDB 5GLH) (Table 1). The datasets for the ET-3 bound receptor were mainly collected with an automated data-collection system, ZOO (K.Y., G.U., K.H., M.Y., and K.H., submitted). This system allowed the convenient collection of a large number of datasets and the determination of the highest-resolution agonist-bound GPCR structures. The electron densities for the agonists in both structures were clearly observed in the *F_o_* − *F_c_* omit maps (Supplementary Fig. 1c, d).

**Table 1.**
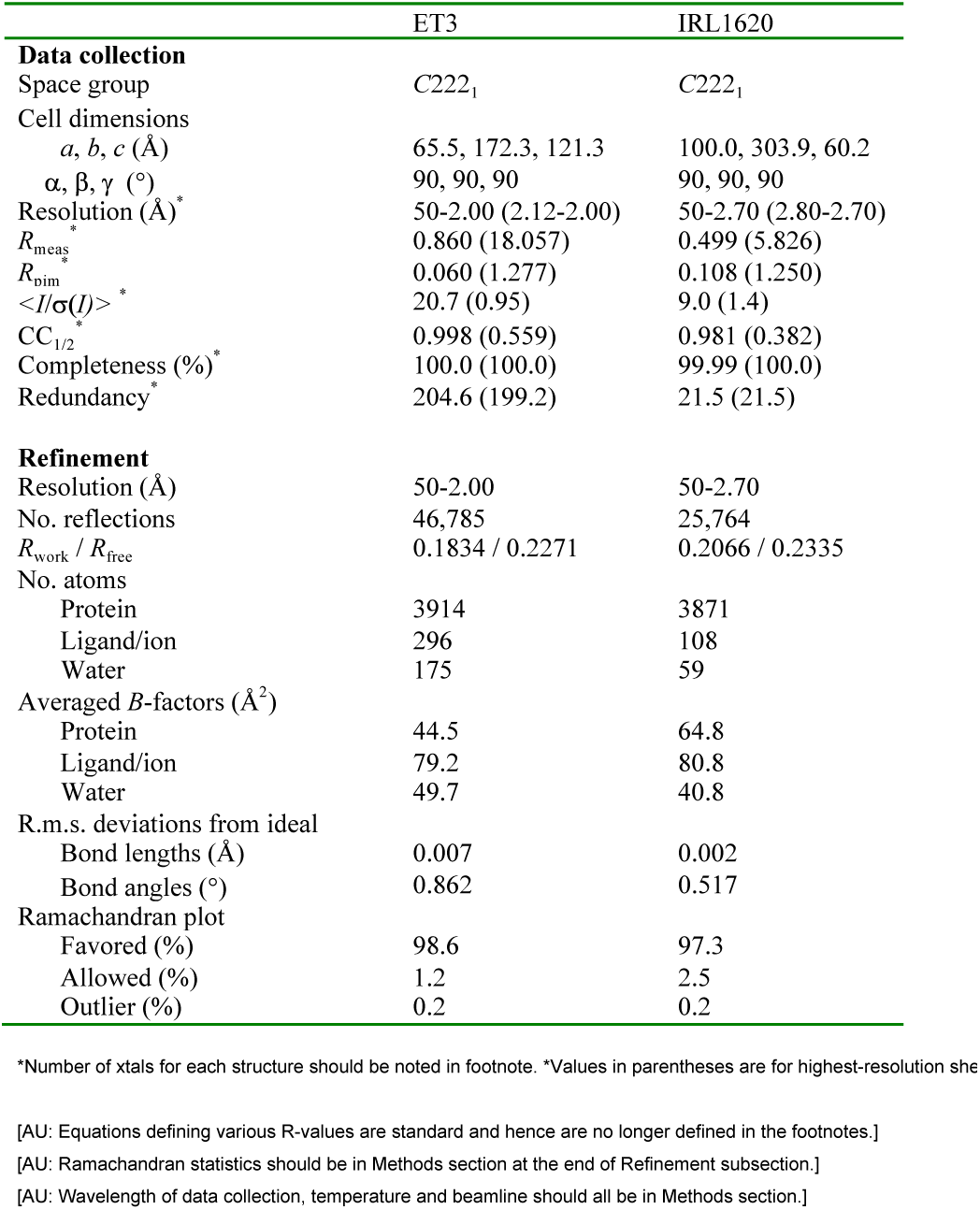
Data collection and refinement statistics.

### ET_B_ structure in complex with the full agonist ET-3

We first describe the ET_B_ structure in complex with ET-3. The overall structure consists of the canonical 7 transmembrane helices (TM), the amphipathic helix 8 at the C-terminus (H8), two antiparallel β-strands in the extracellular loop 2 (ECL2), and the N-terminus that is anchored to TM7 by a disulfide bond (Fig. 2a, b), and is similar to the previous ET-1-bound structure^27^ (overall R.M.S.D of 1.0 Å for the C_α_ atoms) (Fig. 2c). Similar to ET-1, ET-3 adopts a bicyclic architecture comprising the N-terminal region (residues 1-7), the α-helical region (residues 8-17), and the C-terminal region (residues 18-21), and the N-terminal region is attached to the central α-helical region by the intrachain disulfide bond pairs (C1-C15 and C3-C11). The amino acid residues of the α-helical and C-terminal regions are highly conserved between ET-1 and ET-3 (Fig. 2a), and the agonist peptides are superimposed well (Fig. 2b and Supplementary Fig. 2a-d). Accordingly, these regions form similar interactions with the receptor in both structures (Supplementary Fig. 3a, b). In contrast, all of the residues, except for the disulfide bond-forming C1 and C3, are replaced with bulkier residues in ET-3 (Fig. 2a). Despite these sequence differences, the N-terminal regions are similarly accommodated in the orthosteric pocket in both structures, because these bulky residues are exposed to the solvent and poorly interact with the receptor (Fig. 2d). These structural features explain the similar high affinity binding of ET-3 to the ET_B_ receptor, as compared with ET-1.

**Fig. 2.**
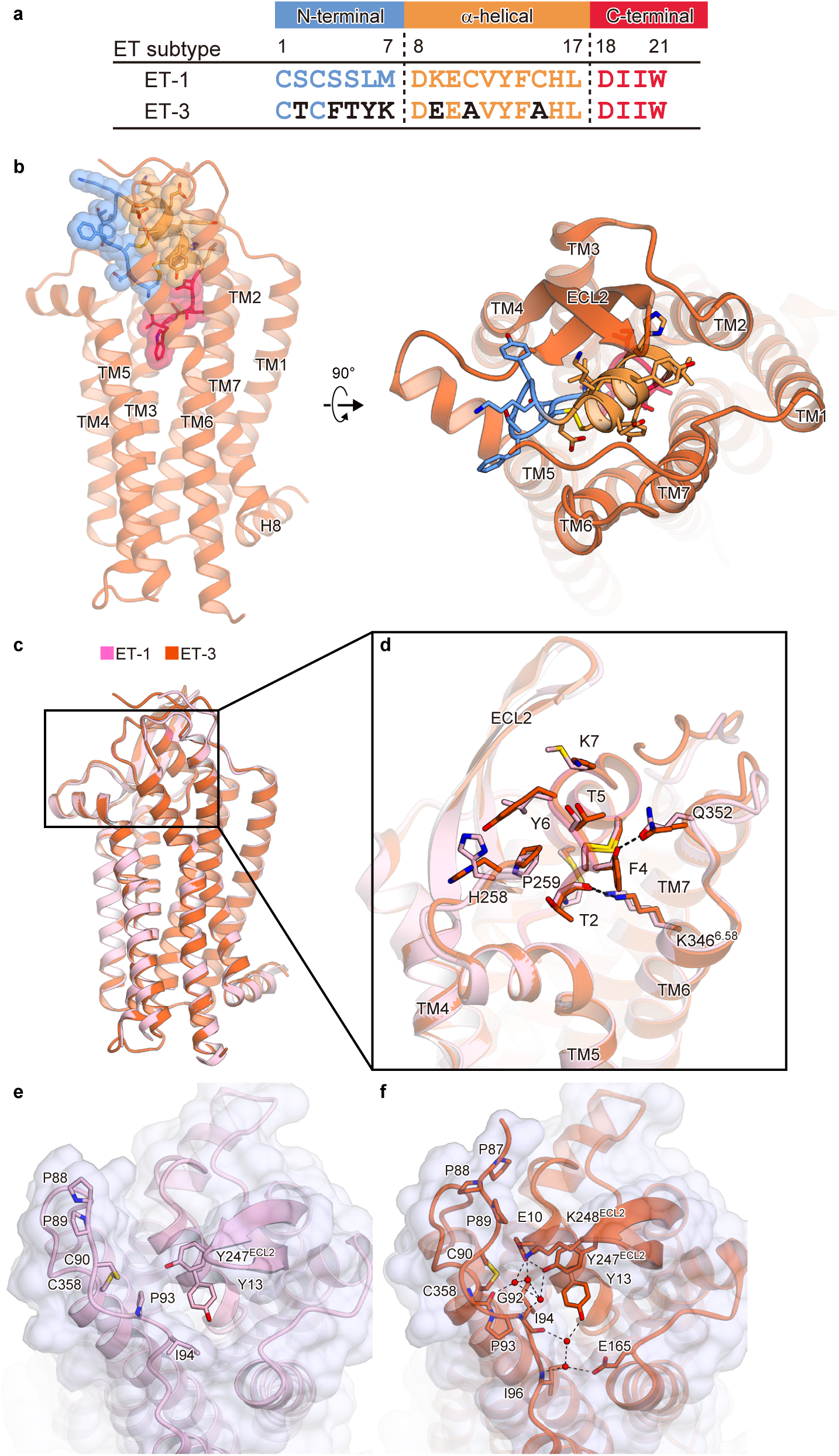
ET_B_ structure in complex with ET-3. **a**, Comparison of the amino acid sequences of ET-1 and ET-3. **b**, The overall structure of the ET-3-bound ET_B_ receptor. The receptor is shown as an orange-red ribbon model. ET-3 is shown as a transparent surface representation and a ribbon model, with its N-terminal region coloured cyan, α-helical region orange, and C-terminal region deep pink. The side chains of ET-3 are shown as sticks, **c**, **d**, Superimposition of the ET-3 and ET-1-bound ET_B_ receptors, coloured orange-red and pink, respectively. Panel (**c**) is viewed from the membrane plane, and panel (**d**) is focused on the N-terminal regions of both structures. The side chains of ET-1 and ET-3 are shown as sticks. **e**, **f**, Comparison of the interactions in the ET-1- (**e**) and ET-3- (**f**) bound structures. The receptors are shown as transparent surface representations and ribbon models. Endothelins are shown as ribbon models. The residues involved in the irreversible binding of the endothelins are shown as sticks.

Previous studies demonstrated that the N-terminal residues of the ET_B_ receptor play a critical role in the virtually irreversible binding of the endothelins^32^. As in the ET-1-bound structure, the N-terminal tail is anchored to TM7 via a disulfide bond between C90 and C358 in the ET-3-bound structure, constituting a lid that prevents agonist dissociation. The high-resolution ET-3-bound structure allowed more accurate tracing of the elongated N-terminal residues (Fig. 2e, f, Supplementary Fig. 1e), as compared with the ET-1-bound structure and revealed more extensive interactions with the agonist peptide. P88, I94, Y247^ECL2^, and K248^ECL2^ form a lid over ET-3, which is stabilized by a water-mediated hydrogen bonding network among the carbonyl oxygen of P93, the side chains of Y247^ECL2^ and K248^ECL2^, and D8 of ET-3 (Fig. 2e). In addition, three consecutive prolines (P87, 88, 89) stretch over the N-terminal region of ET-3. Moreover, ECL1, 2 and the N-terminal residues form an extended water-mediated hydrogen bonding network over ET-3. These extensive interactions strongly prevent the agonist dissociation.

### ET_B_ structure in complex with the partial agonist IRL1620

Next we describe the ET_B_ structure in complex with the partial agonist IRL1620, a linear peptide analogue of ET-1^17^ (Fig. 3a). Previous mutant and structural studies revealed that the N-terminal region contributes to the stability of the overall bicyclic structure by the intramolecular disulfide bonds, and thus facilitates the receptor interaction^17,27,33^. IRL1620 completely lacks the N-terminal region, and consists of only the α-helical and C-terminal regions (Fig. 3b). Two cysteines in the α-helical region are replaced with alanines, and negative charges are introduced into the N-terminal end of the helix, by replacing lysine with glutamic acid (E9) and modifying the N-terminal amide group with a succinyl group (Fig. 3c). The consequent cluster of negative charges on the N-terminal end of IRL1620 (succinyl group, D8, E9, and E10) reportedly plays an essential role in IRL1620 binding to the ET_B_ receptor^17^. This cluster electrostatically complements the positively charged ET_B_ receptor pocket, which includes K346^6.58^ and R357^ECL3^ (Fig. 3d). Moreover, these negative charges neutralize the N-terminal dipole moment of the α-helical region of IRL1620, and thus probably contribute to the stability of the α-helical fold^34^. Due to these interactions, IRL1620 adopts a similar helical conformation, even without the intramolecular disulfide bonds (Supplementary Fig. 2c). IRL1620 thus forms essentially similar interactions with the receptor, as compared with the endogenous agonists, ET-1 and ET-3 (Supplementary Fig. 3a-c). These structural features are consistent with the high affinity of IRL1620 to the ET_B_ receptor, which is comparable to those of ET-1 and ET-3 (Fig. 1).

**Fig. 3.**
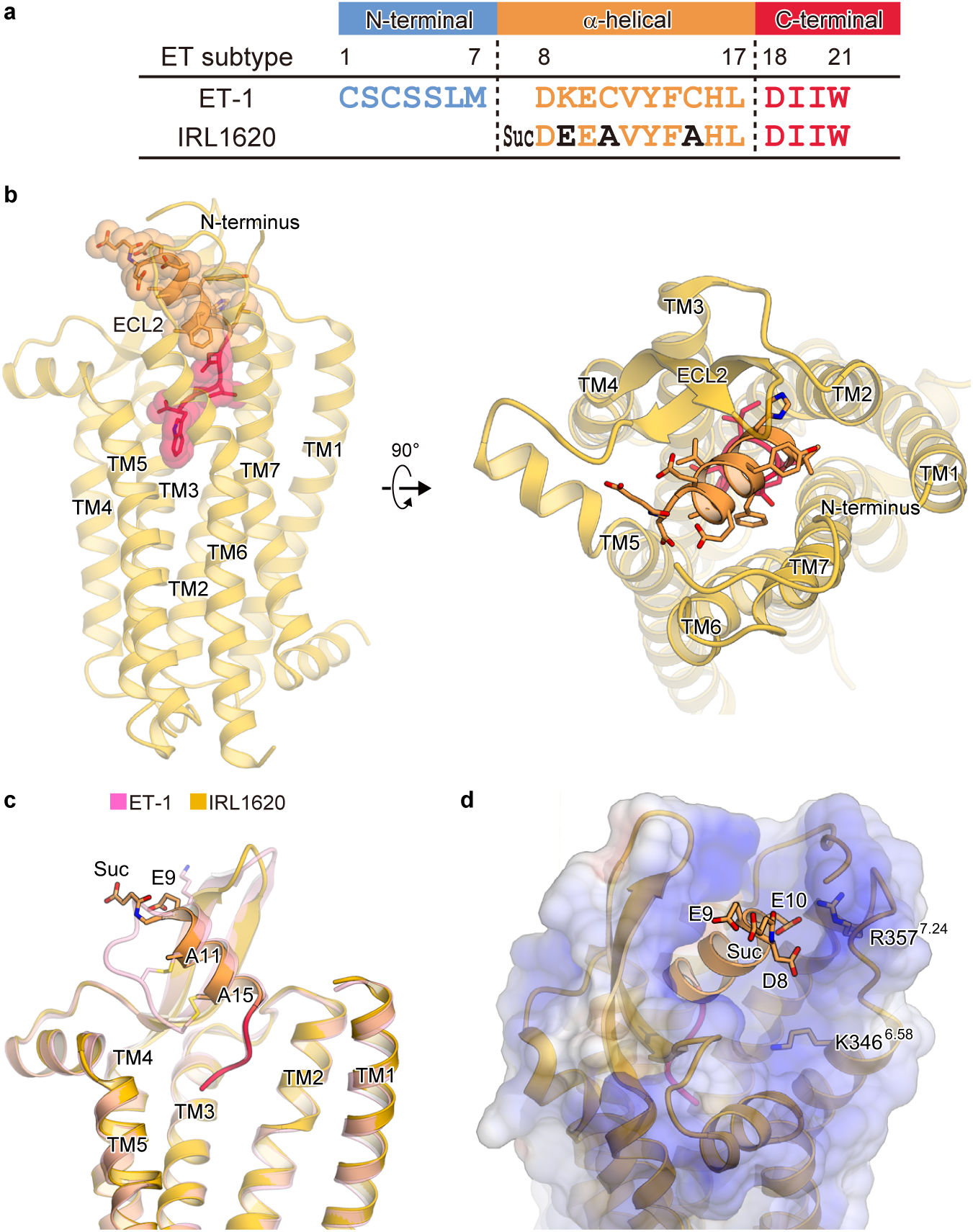
ET_B_ structure in complex with IRL 1620. **a**, Comparison of the amino acid sequences of ET-1 and IRL1620. **b**, The overall structure of the IRL1620-bound ET_B_ receptor. The receptor is shown as an orange ribbon model. IRL1620 is shown as a transparent surface representation and a ribbon model, with the α-helical and C-terminal regions coloured orange and deep pink, respectively. **c**, Superimposition of the IRL1620- and ET-1-bound receptors viewed from the membrane plane, coloured orange and pink, respectively. The ET_B_ receptors and agonists are shown as ribbon models. The different residues between ET-1 and IRL1620 are shown as sticks. **d**, Electrostatic surfaces of the IRL1620-bound ET_B_ structure. The negatively charged moieties on the N-terminal end of IRL1620 and the positively charged residues in the extracellular side of TMs 6 and 7 are shown as sticks.

By contrast, IRL1620 does not bind to the ET_A_ receptor at all in the same concentration range, confirming its high selectivity for the ET_B_ receptor (Fig. 1). To elucidate the mechanism of this selectivity, we compared the amino acid compositions of the IRL1620 binding sites between the ET_B_ and ET_A_ receptors (Fig. 4a and Supplementary Fig. 4). While the transmembrane region is highly conserved, the residues in ECL2 are diverse. In particular, the hydrophobic residues L252 ^ECL2^ and I254 ^ECL2^ are replaced with the polar residues H236 and T238 in the ET_A_ receptor, respectively (Fig. 4a, b). These residues form extensive hydrophobic interactions with the middle part of IRL1620. However, the double mutation of L252H and I254T only reduced the EC_50_ values of IRL1620 by 2-fold in the TGFα shedding and β-arrestin recruitment assays (Fig. 4c), suggesting that these residues are not the sole determinants for the receptor selectivity of IRL1620. Therefore, we focused on other residues of ECL2. In the ET_B_ receptor, P259^ECL2^ and V260^ECL2^ generate a short kink on the loop between the β-strand and TM5, but The ET_A_ receptor has a truncated loop region and completely lacks these residues. (Fig. 4a). In addition, the ET_A_ receptor has a proline (P228) in the first half of ECL2, which should disturb the β-strand formation as in the ET_B_ receptor (Fig. 4b). These observations suggest that ECL2 adopts completely different structures between the ET_A_ and ET_B_ receptors, which may account for their different selectivities. ECL2 of the ET_A_ receptor may form interactions with the N-terminal region of the endothelin peptides, while such interactions are not formed in the ET_B_ receptor, and thus the truncation of the N-terminal region results in the drastically reduced affinity for only the ET_A_ receptor.

**Fig. 4.**
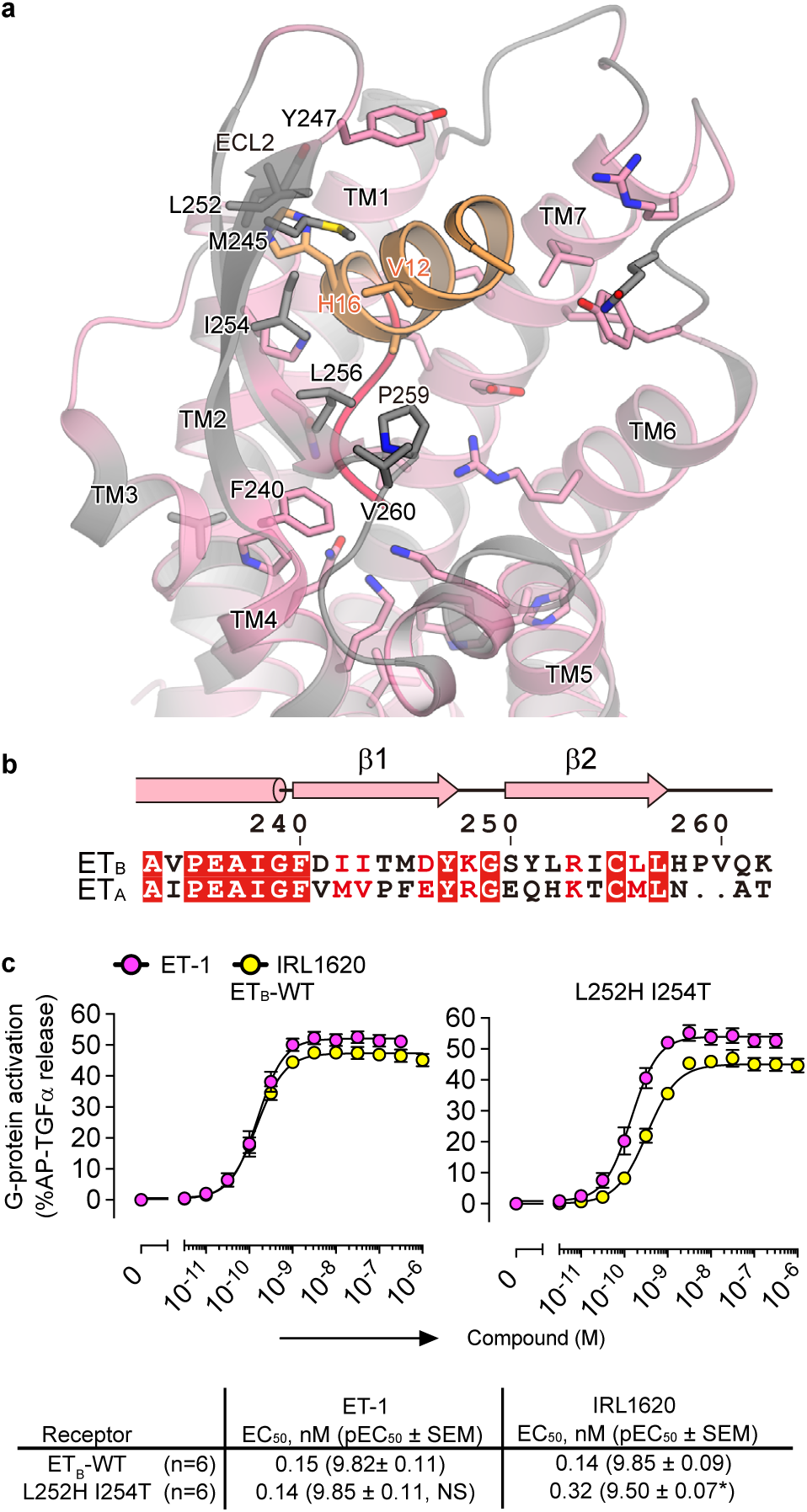
Conservation of the IRL1620 binding site. **a**, Sequence conservation of the IRL1620 binding site between ET_A_ and ET_B_, mapped onto the IRL1620-bound structure. Conserved and non-conserved residues are coloured pink and gray, respectively. The ET_B_ receptor is shown as ribbons, and the residues that form direct interactions with IRL1620 are shown as sticks. **b**, Alignment of the amino acid sequences of the human ET_B_ receptor and human ET_A_ receptor, focused on ECL2. **c**, Dose-response curves of AP-TGFα release response upon IRL1620 treatments of HEK293 cells expressing the wild type receptor and the L252H I254T double mutant. Symbols and error bars are means and s.e.m. (standard error of the mean). The lower panel shows the EC_50_ and E_max_ values. EC_50_ values that significantly differ from the wild-type are denoted by asterisks. \**P* < 0.05, one-way ANOVA with Dunnett’s post hoc test. NS, not significant.

### Structural insight into receptor activation and partial activation

To elucidate the mechanism of the partial activation by IRL1620, we compared the IRL1620-bound structure with the full-agonist ET-3-bound structure (Fig. 5). The intracellular portions of the receptors are quite similar between the ET-3- and IRL1620-bound structures, in which TM7 and H8 adopt active conformations, while the remaining parts of the receptors still represent the inactive conformation of GPCRs (Fig. 5a, Supplementary Fig. 5). On the extracellular side, IRL1620 induces similar conformational changes to those observed in the ET-1 and ET-3 structures; namely, the large inward motions of TM2, 6, and 7, which are critical for receptor activation (Fig. 5b, c). However, the extent of the inward motions of TM6-7 is smaller by about 1 Å in the IRL1620-bound structure, as compared with the ET-3-bound structure, due to the different ligand architectures between IRL1620 and ET-3. Since IRL1620 lacks the N-terminal region, the orthosteric pocket of the receptor has more space, and consequently the α-helical region of IRL1620 is tilted differently toward TM6 (Fig. 5b). In addition, while the N-terminal region of ET-3 interacts with TM6 of the receptor, by forming a hydrogen bond between the carbonyl oxygen of T2 and K346^6.58^ (superscripts indicate Ballesteros–Weinstein numbers), IRL1620 lacks this interaction, resulting in the different orientation of TM6-7. As TM6 plays an especially important role for the cytoplasmic G-protein binding, this difference is probably related to the partial agonist activity of IRL1620.

**Fig. 5.**
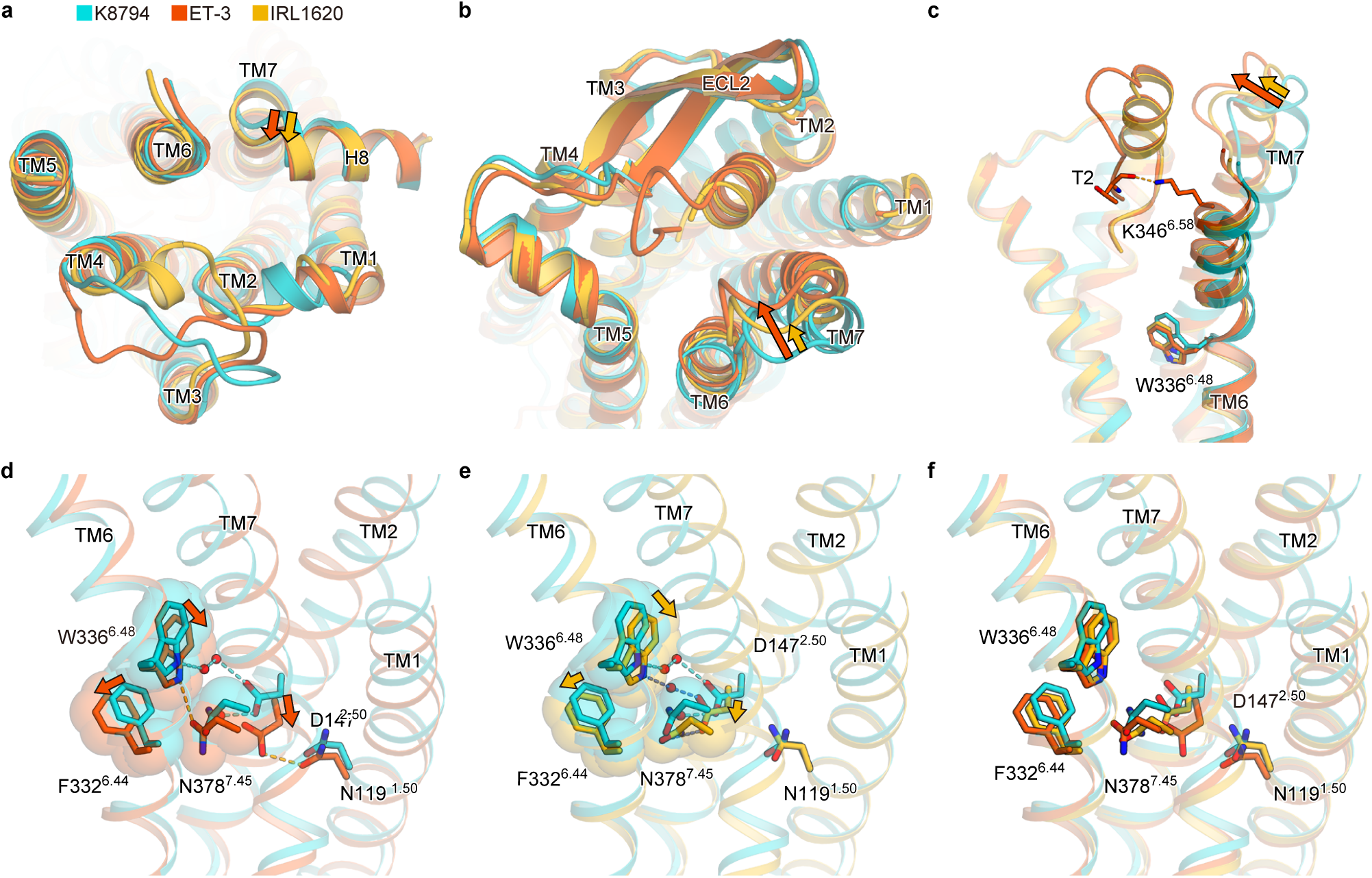
Comparsion of the K8794, ET-3, and IRL1620-bound structures. **a-c**, Comparison of the K-8794-bound inactive, IRL1620-bound partially active, and ET-3-bound active ET_B_ structures, coloured turquoise, orange, and orange-red, respectively. The ET_B_ receptors and endothelins are shown as ribbon models. (**a**), (**b**), and (**c**) are viewed from the intracellular side, the extracellular side, and the membrane plane, respectively. The cytoplasmic cavity for the G-protein binding is still hindered by the inwardly-oriented TM6 in the ET-3 and IRL1620-bound receptors, representing the inactive conformations. This is consistent with the notion that the fully-active conformation is only stabilized when the G-protein is bound, as shown in the previous nuclear magnetic resonance (NMR) and double electron electron resonance (DEER) spectroscopy studies of GPCRs. **d-f**, Superimposition of the ET_B_ structures bound to K-8794 and ET-3 (**d**), K-8794 and IRL1620 (**e**), and K-8794, IRL1620, and ET-3 (**f**), focused on the intermembrane parts. The receptors are shown as ribbons, and the side chains of N119^1.50^, D147^2.50^, F332^6.44^, W336^6.48^, and N378^7.45^ are shown as sticks with transparent surface representations. Waters are shown as red spheres. The dashed lines show hydrogen bonds coloured in the respective structures.

A comparison of the intermembrane parts revealed further differences in the allosteric coupling between the orthosteric pocket and the intermembrane part. Previous studies have shown that the agonist binding induces the disruption of the hydrogen-bonding network around D147^2.50^, which connects TMs 2, 3, 6, and 7 and stabilizes the inactive conformation of the ET_B_ receptor^28^. The present high resolution ET-3-bound structure provides a precise mechanistic understanding of this rearrangement (Supplementary Fig. 6a–c). In particular, the water-mediated hydrogen bonds involving D147^2.50^, W336^6.48^, and N378^7.45^ in the inactive conformation collapse upon ET-3 binding, by the inward motions of TMs 2, 6, and 7 (Fig. 5d). The W336^6.48^ side chain moves downward by about 2.5 Å, resulting in the disruption of the water-mediated hydrogen bond with D147^2.50^, and consequently, the D147^2.50^ side chain moves downward by about 3 Å and forms hydrogen bonds with the N382^7.49^ and N119^1.50^ side chains. The N378^7.45^ side chain also moves downward by about 1.5 Å and forms a hydrogen bond with the nitrogen atom of the W336^6.48^ side chain. The downward movements of the W336^6.48^ and N378^7.45^ side chains consequently induce the outward repositioning of the F332^6.44^ side chain, by about 1 Å. W6.48 and F6.44 are considered to be the “transmission switch” of the class A GPCRs^35^, which transmits the agonist-induced motions to the cytoplasmic G-protein coupling interface. Overall, our results show that the collapse of the water-mediated hydrogen-bonding network involving D147^2.50^, W336^6.48^, and N378^7.45^ propagates as the structural change in the transmission switch, and probably induces the outward displacement of the cytoplasmic portion of TM6 upon G-protein activation (Fig. 6, Supplementary Fig. 5).

**Fig. 6.**
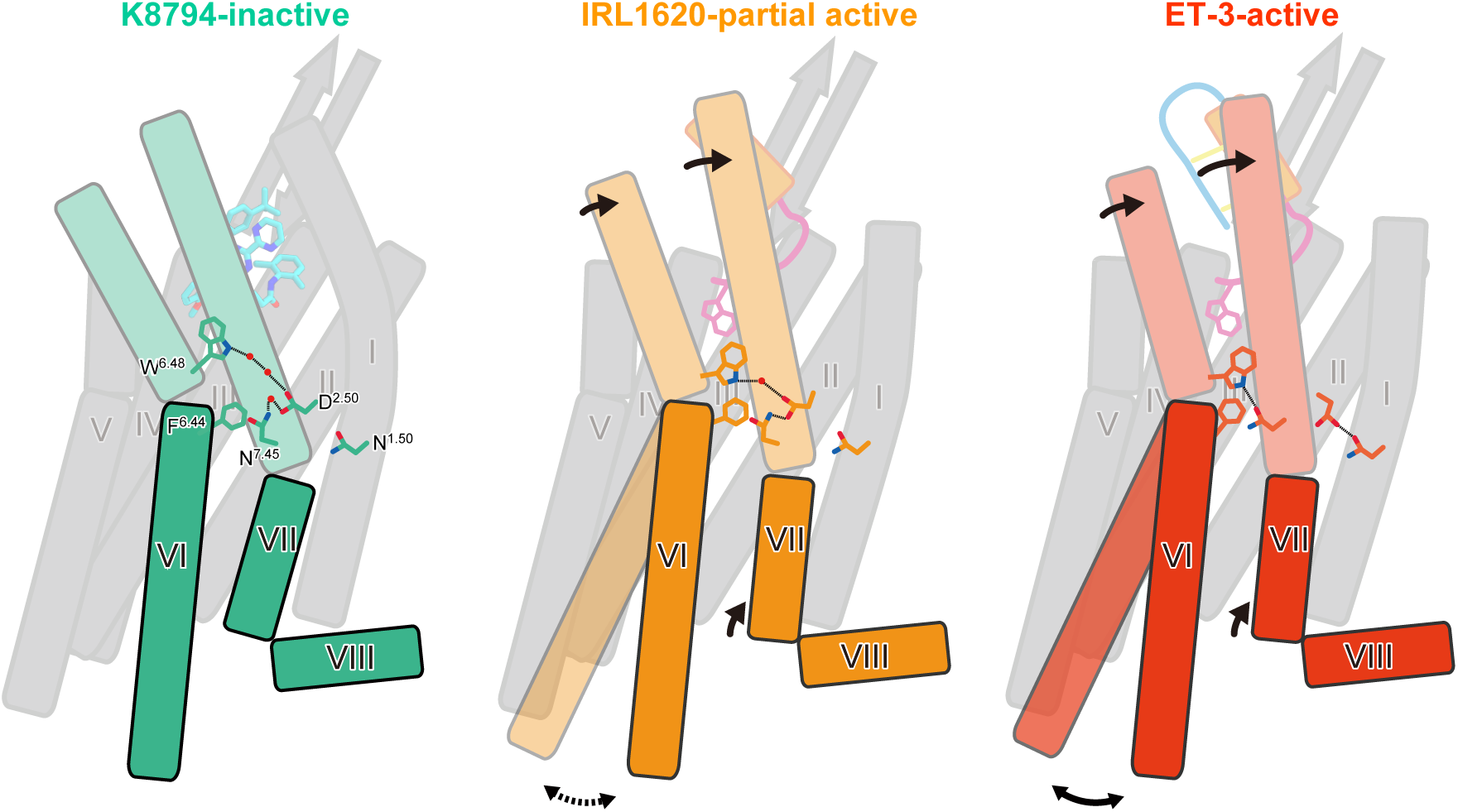
Schematic representations of receptor activation by ET-3 and partial activation by IRL 1620. Schematic representations of receptor activation by ET-3 and partial activation by IRL1620. TM6, TM7, and H8 are highlighted. The residues involved in the signal transduction (N1.50, D2.50, F6.44, W6.48, and N7.45) are represented with sticks. Hydrogen bonds are indicated by black dashed lines. Black arrows indicate the conformational changes in TM6 and TM7 upon ET-3 and IRL1620 binding. In the K-8794-bound structure (left), a water-mediated hydrogen bonding network among D147^2.50^, W336^6.48^, and N378^7.45^ stabilizes the inactive conformation of the receptor. ET-3 binding disrupts this network and propagates the structural change of the transmission switch comprising F332^6.44^ and W336^6.48^ (middle), leading to the outward movement of TM6 upon G-protein coupling. In the IRL1620-bound structure (right), the water-mediated hydrogen-binding network is preserved, thus preventing the outward movement of TM6 upon G-protein coupling.

IRL1620 induces a similar but slightly different rearrangement of the hydrogen bonding network in the intermembrane part (Fig. 5e). Due to the smaller inward shift of TM6, the downward shift of the W336^6.48^ side chain is smaller in the IRL1620-bound structure, and it still forms a water-mediated hydrogen bond with the D147^2.50^ side chain. Consequently, the D147^2.50^ side chain forms a direct hydrogen bond with N378^7.45^, thereby preventing the downward motion of N378^7.45^ and the hydrogen bond formation between N378^7.45^ and W336^6.48^. Overall, the downward motions of the W336^6.48^ and N378^7.45^ side chains are only moderate, as compared to those in the ET-3-bound structure, and the hydrogen-bonding network involving D147^2.50^, W336^6.48^, and N378^7.45^ is partially preserved in the IRL1620-bound structure (Fig. 5e, Supplementary Fig. 6). Accordingly, in the IRL1620-bound structure, the position of the “transmission switch” residue F332^6.44^ is in between those of the active (ET-3 bound) and inactive (K-8794-bound) structures (Fig. 5f). This intermediate position of F332^6.44^ should partly prevent the outward displacement of the cytoplasmic portion of TM6 that is required for G-protein activation, thus accounting for the partial agonistic effect of IRL1620 (Fig. 6).

## Discussion

Previous studies have suggested that the α-helical and C-terminal regions of endothelins are critical elements for receptor activation^36-38^, while the N-terminal region is only responsible for the ETR selectivity^6^. Indeed, the N-terminal region-truncated analog IRL1620 has similar EC_50_ values, as compared with ET-1^17^. However, our pharmacological experiments for the first time proved that IRL1620 functions as a partial agonist for the ET_B_ receptor, rather than a full agonist, suggesting the participation of the N-terminal region in the activation process of the ET_B_ receptor. To clarify the receptor activation mechanism, we determined the crystal structures of the human ET_B_ receptor in complex with ET-3 and IRL1620. The high-resolution structure of the ET-3-bound ET_B_ receptor revealed that the large inward motions of the extracellular portions of TMs 2, 6, and 7 disrupt the water-mediated hydrogen bonding network at the receptor core, leading to receptor activation. The IRL1620-bound ET_B_ structure revealed that the IRL1620-induced inward motions of TMs 6 and 7 are smaller by about 1 Å, as compared with those caused by ET-3, due to the lack of the N-terminal region. Consequently, the hydrogen-bonding network at the receptor core is partially preserved, thus preventing the transition to the fully active conformation adopted upon G-protein binding. These observations suggest that the interactions between the N-terminal regions of endothelins and TM6 also participate in receptor activation, while the extensive interactions of the α-helical and C-terminal regions with the receptor primarily contribute to this process (Supplementary Fig. 3c). This activation mechanism is different from that of the small-molecule activated GPCRs (*e.g*., β2 adrenaline and M2 muscarinic acetylcholine receptors), in which only a small number of hydrogen-bonding interactions between the agonist and the receptor induce receptor activation, by affecting the receptor dynamics^39,40^.

D2.50 is one of the most conserved residues among the class A GPCRs (90%). Recent high-resolution structures have revealed that a sodium ion coordinates with D2.50 and forms a water-mediated hydrogen bonding network in the intermembrane region, which stabilizes the inactive conformation of the receptor^41^, and its collapse leads to receptor activation. Our previous 2.2 Å resolution structure of the K-8794-bound ET_B_ receptor revealed that a water molecule occupies this allosteric sodium site and participates in the extensive hydrogen-bonding network, instead of a sodium ion^28^, and this hydrogen-bonding network is collapsed in the 2.8 Å resolution structure of the ET-1-bound ET_B_ receptor, indicating its involvement in the receptor activation^27^. Nevertheless, the precise rearrangement of this network still remained to be elucidated, due to the limited resolution. The current 2.0 Å resolution structure of the ET-3-bound ET_B_ receptor revealed that the collapse of the water-mediated interaction between W336^6.48^ and D147^2.50^ is critical for receptor activation. This network is still partly preserved in the IRL1620-bound structure, thus preventing the transition to the fully active conformation upon G-protein coupling (Fig. 6). W6.48 is also highly conserved among the class A GPCRs (71%)^42^, and the association between W6.48 and D2.50 plays a critical role in the GPCR activation process, as shown in the previous nuclear magnetic resonance (NMR) study of the adenosine A_2A_ receptor^43^. Given the importance of W3.36 and D2.50 in the activation of GPCRs, our proposed model of the partial receptor activation by IRL1620 is consistent with the previous functional analyses of GPCRs. To date, the β_1_ adrenergic receptor is the only receptor for which agonist- and partial agonist-bound structures were reported^44^. However, these structures are both stabilized in inactive conformations by the thermostabilizing mutations and thus revealed only slight differences (Supplementary Fig. 7). Therefore, our study provides the first structural insights into the partial activation of class A GPCRs.

Our current study further suggests possible improvements in clinical studies using ET_B_-selective agonists. IRL1620 is the smallest among the ET_B_-selective agonists, and thus is expected to be useful for the treatment of cancers and other diseases^18–22,24,25^. While its effectiveness has been proven in rat experiments, a recent phase 2 study has failed^26^, and thus further improvement of IRL1620 is required for clinical applications. Our cell-based assays and structural analysis revealed the partial agonistic effect of IRL1620 on the ET_B_ receptor, suggesting the possible enhancement of its efficacy. The development of an ET_B_-selective full agonist based on the IRL1620-bound ET_B_ structure will be beneficial for clinical applications.

## Materials and methods

### Expression and purification

The haemagglutinin signal peptide, followed by the Flag epitope tag (DYKDDDDK) and a nine-amino-acid linker, was added to the N-terminus of the receptor, and a tobacco etch virus (TEV) protease recognition sequence was introduced between G57 and L66, to remove the disordered N terminus during the purification process. The C-terminus was truncated after S407, and three cysteine residues were mutated to alanine (C396A, C400A, and C405A) to avoid heterogeneous palmitoylation. To improve crystallogenesis, T4 lysozyme containing the C54T and C97A mutations was introduced into intracellular loop 3, between L303^5.68^ and L311^6.23^ (ET_B_-Y5-T4L).

The thermostabilized construct ET_B_-Y5-T4L was subcloned into a modified pFastBac vector, with the resulting construct encoding a TEV cleavage site followed by a GFP-His^10^ tag at the C-terminus. The recombinant baculovirus was prepared using the Bac-to-Bac baculovirus expression system (Invitrogen). Sf9 insect cells were infected with the virus at a cell density of 4.0 × 10^6^ cells per millilitre in Sf900 II medium, and grown for 48 h at 27 °C. The harvested cells were disrupted by sonication, in buffer containing 20 mM Tris-HCl, pH 7.5, and 20% glycerol. The crude membrane fraction was collected by ultracentrifugation at 180,000*g* for 1 h. The membrane fraction was solubilized in buffer, containing 20 mM Tris-HCl, pH 7.5, 200 mM NaCl, 1% DDM, 0.2% cholesterol hemisuccinate, and 2 mg ml^−1^ iodoacetamide, for 1 h at 4 °C. The supernatant was separated from the insoluble material by ultracentrifugation at 180,000*g* for 20 min, and incubated with TALON resin (Clontech) for 30 min. The resin was washed with ten column volumes of buffer, containing 20 mM Tris-HCl, pH 7.5, 500 mM NaCl, 0.1% LMNG, 0.01% CHS, and 15 mM imidazole. The receptor was eluted in buffer, containing 20 mM Tris-HCl, pH 7.5, 500 mM NaCl, 0.01% LMNG, 0.001% CHS, and 200 mM imidazole. The eluate was treated with TEV protease and dialysed against buffer (20 mM Tris-HCl, pH 7.5, and 500 mM NaCl). The cleaved GFP–His_10_ tag and the TEV protease were removed with Co^2+^-NTA resin. The receptor was concentrated and loaded onto a Superdex200 10/300 Increase size-exclusion column, equilibrated in buffer containing 20 mM Tris-HCl, pH 7.5, 150 mM NaCl, 0.01% LMNG, and 0.001% CHS. Peak fractions were pooled, concentrated to 40 mg ml^−1^ using a centrifugal filter device (Millipore 50 kDa MW cutoff), and frozen until crystallization. During the concentration, ET-3 or IRL1620 was added to a final concentration of 100 μM.

### Crystallization

The purified receptors were reconstituted into the lipidic cubic phase (LCP) of monoolein (Nucheck), supplemented with cholesterol at a ratio of 8:9:1 (w/w) for protein:monoolein-cholesterol. The protein-laden mesophase was dispensed into 96-well glass plates in 30 nl drops and overlaid with 800 nl precipitant solution, using a mosquito LCP (TTP LabTech)^45,46^. Crystals of ET_B_-Y5-T4L bound to ET-3 were grown at 20°C in the precipitant conditions containing 25% PEG500DME, 100 mM MES-N_a_OH, pH 6.0, and 100 mM ammonium citrate tribasic. The crystals of ET_B_-Y5-T4L bound to IRL1620 were grown in the precipitant conditions containing 20-25% PEG500DME, 100 mM sodium citrate, pH 5.0, and 100 mM (NH_4_)_2_SO_4_ or NH_4_Cl. The crystals were harvested directly from the LCP using micromounts (MiTeGen) or LithoLoops (Protein Wave) and frozen in liquid nitrogen, without adding any extra cryoprotectant.

### Data collection and structure determination

X-ray diffraction data were collected at the SPring-8 beamline BL32XU with 1×10 to 8 × 25 μm^2^ (width × height) micro-focused beams and an EIGER X 9M detector (Dectris). For the IRL1620 data, we manually collected 68 data sets (10° to 180° per crystal), and the collected images were automatically processed with KAMO^31^ (https://github.com/keitaroyam/yamtbx). Each data set was indexed and integrated with XDS^47^ and then subjected to a hierarchical clustering analysis based on the unit cell parameters using BLEND^48^. After the rejection of outliers, 46 data sets were finally merged with XSCALE^47^. From the ET-3-bound crystals, various wedge data sets (3-180°) per crystal were mainly collected with the ZOO system, an automatic data-collection system developed at SPring-8 (K.Y., G.U., K.H., M.Y., and K.H., submitted). The loop-harvested microcrystals were identified by raster scanning and subsequently analyzed by SHIKA^49^. The collected images were processed in the same manner, except that correlation coefficient-based clustering was used instead of BLEND, and finally 483 datasets were merged. The ET-3-bound structure was determined by molecular replacement with PHASER^50^, using the ET-1-bound ET_B_ structure (PDB 5GLH). Subsequently, the model was rebuilt and refined using COOT^51^ and PHENIX^52^, respectively. The IRL1620-bound structure was determined by molecular replacement, using the ET-1-bound ET_B_ structure, and subsequently rebuilt and refined as described above. The final model of ET-3-bound ET_B_-Y5-T4L contained residues 86-303 and 311-403 of ET_B_, all residues of T4L, ET-3, 12 monoolein molecules, a citric acid, and 175 water molecules. The final model of IRL1620-bound ET_B_-Y5-T4L contained residues 87–303 and 311-402 of ET_B_, all residues of T4L, IRL1620, 3 monoolein molecules, 4 sulfate ions, a citric acid, and 59 water molecules. The model quality was assessed by MolProbity^53^. Figures were prepared using CueMol (http://www.cuemol.org/ja/)

### TGFα shedding assay

The TGFα shedding assay, which measures the activation of Gq and G12 signaling^47^, was performed as described previously^28^. Briefly, a plasmid encoding an ET_B_ construct with an internal FLAG epitope tag or an ET_A_ construct was transfected, together with a plasmid encoding alkaline phosphatase (AP)-tagged TGFα (AP-TGFα), into HEK293A cells by using a polyethylenimine (PEI) transfection reagent (1 µg ETR plasmid, 2.5 µg AP-TGFα plasmid and 25 µl of 1 mg/ml PEI solution per 10-cm culture dish). After a one day culture, the transfected cells were harvested by trypsinization, washed, and resuspended in 30 ml of Hank’s Balanced Salt Solution (HBSS) containing 5 mM HEPES (pH 7.4). The cell suspension was seeded in a 96 well plate (cell plate) at a volume of 90 μl per well and incubated for 30 min in a CO_2_ incubator. Test compounds, diluted in 0.01% bovine serum albumin (BSA) and HEPES-containing HBSS, were added to the cell plate at a volume of 10 µl per well. After a 1 h incubation in the CO_2_ incubator, the conditioned media (80 μl) was transferred to an empty 96-well plate (conditioned media (CM) plate). The AP reaction solution (10 mM p-nitrophenylphosphate (p-NPP), 120 mM Tris–HCl (pH 9.5), 40 mM NaCl, and 10 mM MgCl_2_) was dispensed into the cell plates and the CM plates (80 µl per well). The absorbance at 405 nm (Abs405) of the plates was measured, using a microplate reader (SpectraMax 340 PC384, Molecular Devices), before and after a 1 h incubation at room temperature. AP-TGFα release was calculated as described previously^28^. The AP-TGFα release signals were fitted to a four-parameter sigmoidal concentration-response curve, using the Prism 7 software (GraphPad Prism), and the pEC_50_ (equal to -Log_10_ EC_50_) and E_max_ values were obtained.

### β-Arrestin Recruitment Assay

For the NanoBiT-β-arrestin recruitment assay^54^, a receptor construct was designed to fuse the small fragment (SmBiT) of the NanoBiT complementation luciferase to the C-terminus of the ETR construct with a 15-amino acid flexible linker (GGSGGGGSGGSSSGG). A PCR-amplified ETR fragment and an oligonucleotide-synthesized SmBiT were assembled and inserted into the pCAGGS mammalian expression plasmid (a kind gift from Dr. Jun-ichi Miyazaki, Osaka University), using a NEBuilder HiFi DNA Assembly system (New England Biolabs). A β-arrestin construct was generated by fusing the large fragment (LgBiT), with nucleotide sequences gene-synthesized with mammalian codon optimization (Genscript), to the N-terminus of human β-arrestin1 (βarr1) with the 15-amino acid linker. The R393E and R395E mutations, which were shown to abrogate AP-2 binding and thus are defective in internalization^55^, were introduced into LgBiT-βarr1 to facilitate the formation of the ETR-βarr1 complex. The plasmid encoding the ET_B_-SmBiT construct or the ET_A_-SmBiT construct was transfected, together with the plasmid encoding the internalization-defective LgBiT-βarr1, into HEK293A cells by the PEI method (1 µg ETR plasmid, 0.5 µg LgBiT-βarr1 plasmid, and 25 µl of 1 mg/ml PEI solution per 10-cm culture dish). After a one day culture, the transfected cells were harvested with EDTA-containing Dulbecco’s phosphate-buffered saline and resuspended in 10 ml of HBSS containing 5 mM HEPES and 0.01% BSA (BSA-HBSS). The cell suspension was seeded in a 96-well white plate at a volume of 80 μl per well and loaded with 20 µl of 50 µM coelenterazine (Carbosynth), diluted in BSA-HBSS. After an incubation at room temperature for 2 hours, the background luminescent signals were measured using a luminescent microplate reader (SpectraMax L, Molecular Devices). Test compounds (6X, diluted in BSA-HBSS) were manually added to the cells (20 µl). After ligand addition, the luminescent signals were measured for 5 min at 20 sec intervals. The luminescent signal was normalized to the initial count, and the fold-change values over 5–10 min after ligand stimulation were averaged. The fold-change βarr recruitment signals were fitted to a four-parameter sigmoidal concentration-response, and the pEC_50_ and E_max_ values were obtained as described above.

## Acknowledgements

We thank the members of the Nureki lab and the beamline staff at BL32XU of SPring-8 (Sayo, Japan) for technical assistance during data collection. The diffraction experiments were performed at SPring-8 BL32XU (proposal 2016A2527). This work was supported by JSPS KAKENHI grants 16H06294 (O.N.), 17J30010 (W.S.), 30809421 (W.S.), 15H06862 (K.Y.), 17H05000 (T.N.), and 17K08264 (A.I.), and the Core Research for Evolutional Science, PRESTO from the Japan Science and Technology (JST) Technology Program; the Platform for Drug Discovery, Information, and Structural Life Science from the Ministry of Education, Culture, Sports, Science, and Technology of Japan; and the Japan Agency for Medical Research and Development (AMED) grants the PRIME JP17gm5910013 (A.I.) and the LEAP JP17gm0010004 (A.I. and J.A.), and the National Institute of Biomedical Innovation.

## Author contributions

W.S. designed all of the experiments, purified the ET_B_ receptor in complex with ET-3, collected data, and refined the structures. T.I. expressed, purified, and crystallized the ET_B_ receptor in complex with ET-3 and IRL1620, collected data, and refined the structures. A.I., F.M., N.K., and J.A. performed and oversaw the cell-based assays. K.Y. and K.H. developed a pipeline for data collection and processing, and assisted with the structure determination and refinement. The manuscript was prepared by W.S., T.I., A.I., K.Y., T.N., and O.N. T.N. and O.N. supervised the research. The authors declare no competing financial interests. Coordinates and structure factors have been deposited in the Protein Data Bank, under accession numbers XXXX for the ET-3-bound and YYYY for the IRL1620-bound structures. The X-ray diffraction images are also available at SBGrid Data Bank (https://data.sbgrid.org/), under IDs XXX and XXX (ET-3-bound and IRL1620-bound, respectively).

## Supplementary Figure Legends

**Supplementary Fig. 1.**
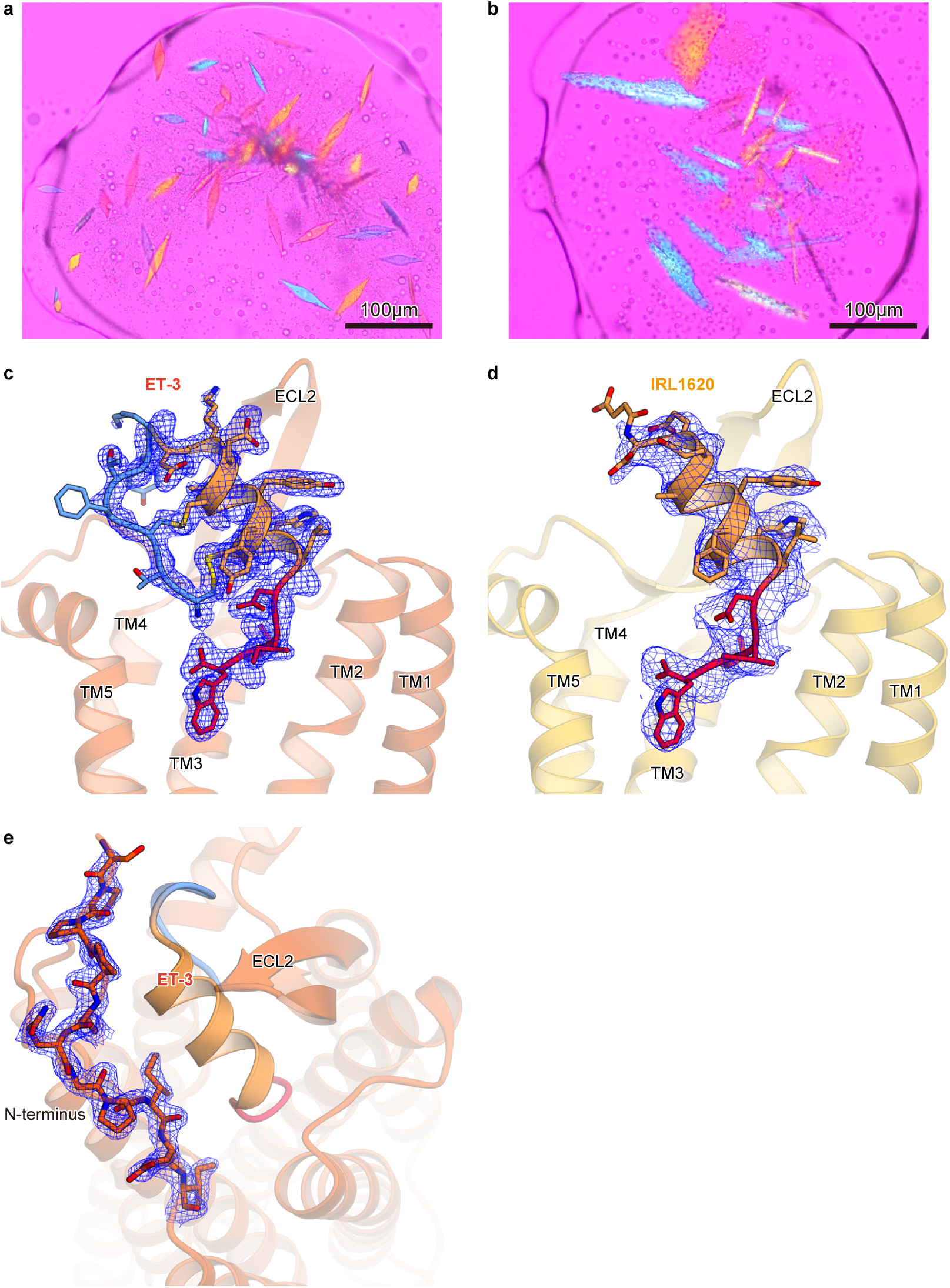
Crystals and electron densities. **a**, **b**, Crystals of the ET_B_ receptors in complex with ET-3 (**a**) and IRL1620 (**b**). **c**, **d**, *F_o_* − *F_c_* omit maps for ET-3 (**c**) and IRL1620 (**d**), contoured at 2.5σ and 1.5σ, respectively. **e**, 2*F_o_* − *F_c_* map of the N-terminus in the ET-3-bound structures, contoured at 2.5σ.

**Supplementary Fig. 2.**
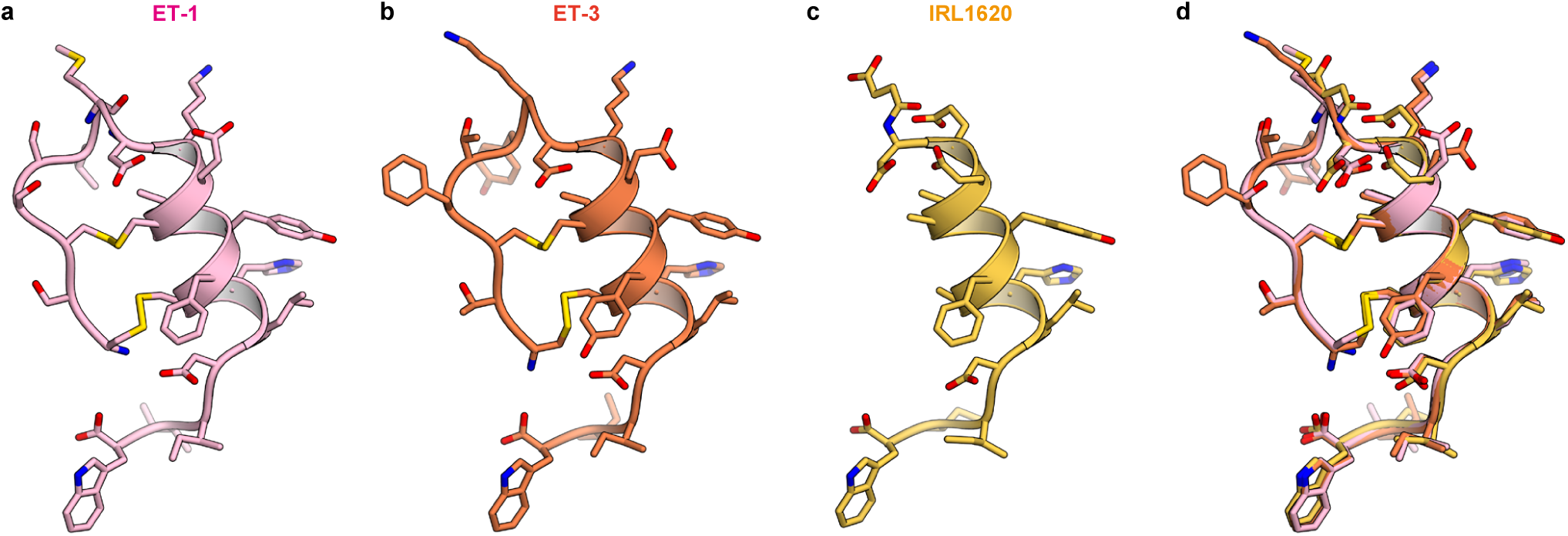
Comparison of agonist structures. **a-c**, Structures of ET-1 (**a**), ET-3 (**b**), and IRL1620 (**c**) in the complex structures, coloured pink, orange-red, and orange, respectively. The agonists are shown as ribbon and stick models. **d**, Superimposition of the endothelin structures.

**Supplementary Fig. 3.**
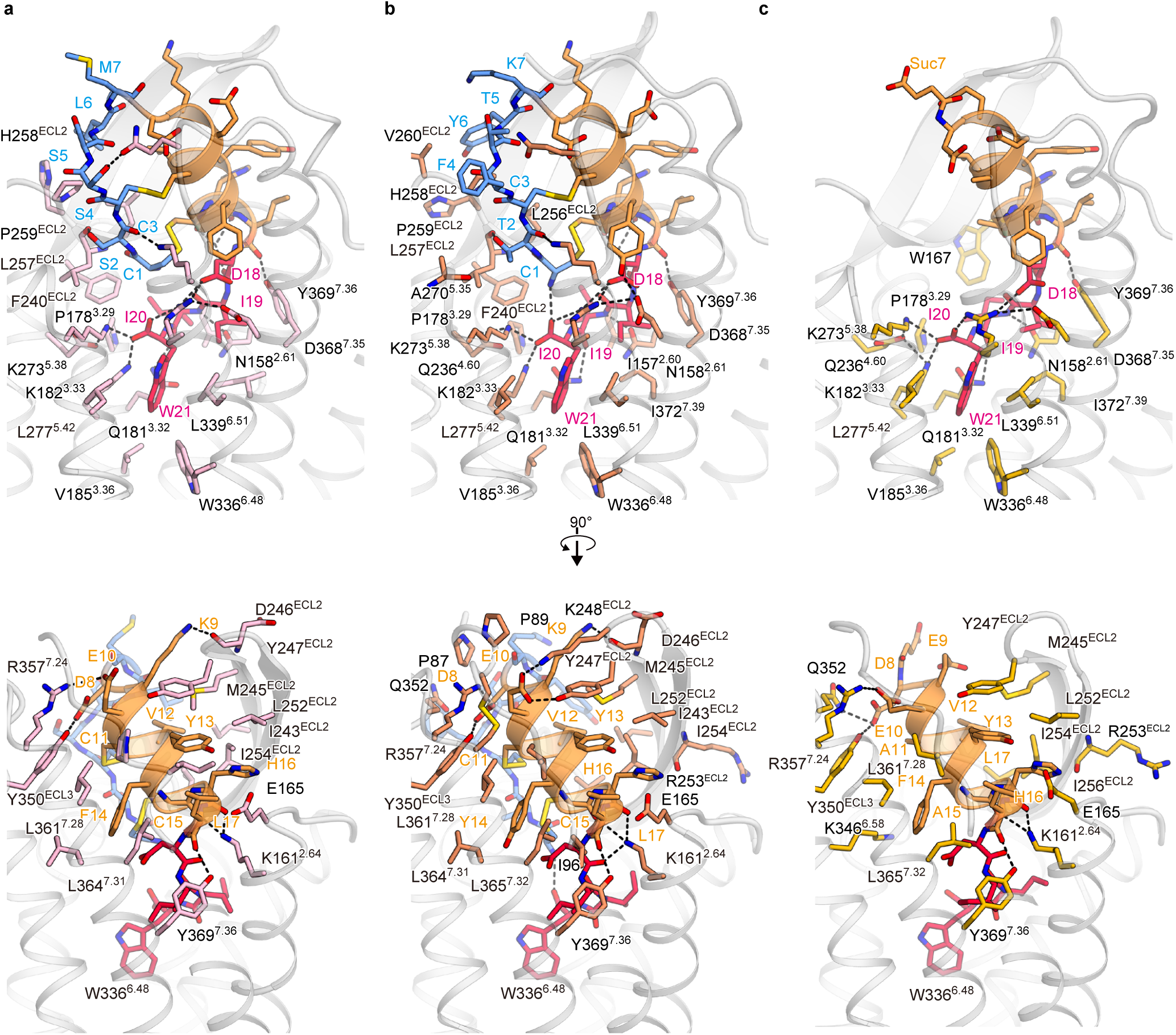
Comparison of receptor interactions of agonist peptides. **a**-**c**, Comparison of the receptor binding interactions of ET-1 (**a**), ET-3 (**b**), and IRL1620 (**c**). The upper panels focus on the interactions of the N-terminal and C-terminal regions, and the lower panels focus on those of the α-helical regions. The agonists are shown as ribbon and stick models, coloured as in Figs. 2b and 3b. The structures of the receptors are shown as silver ribbons. The residues that interact with the agonists are shown as sticks, coloured as in Supplementary Fig. 2.

**Supplementary Fig. 4.**
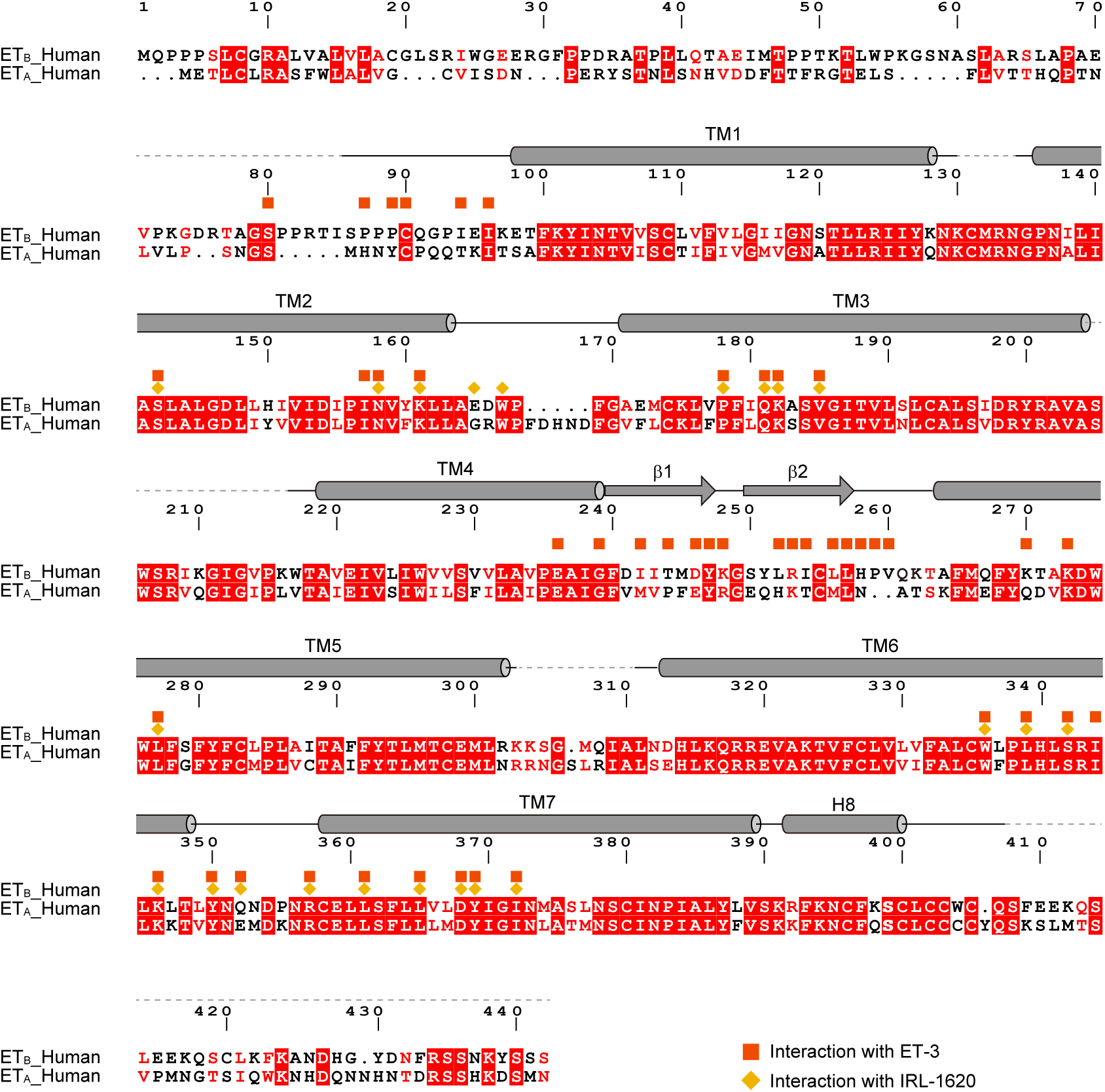
Alignment of human ET_A_ and ET_B_ receptor sequences. Alignment of the amino acid sequences of the human ET_B_ receptor (UniProt ID: P24530) and human ET_A_ receptor (P25101). Secondary structure elements for α-helices and β-strands are indicated by cylinders and arrows, respectively. Conservation of the residues between ET_A_ and ET_B_ is indicated as follows: red panels for completely conserved; red letters for partially conserved; and black letters for not conserved. The residues involved in the ET-3 and IRL1620 binding are indicated with squares and diamonds, respectively.

**Supplementary Fig. 5.**
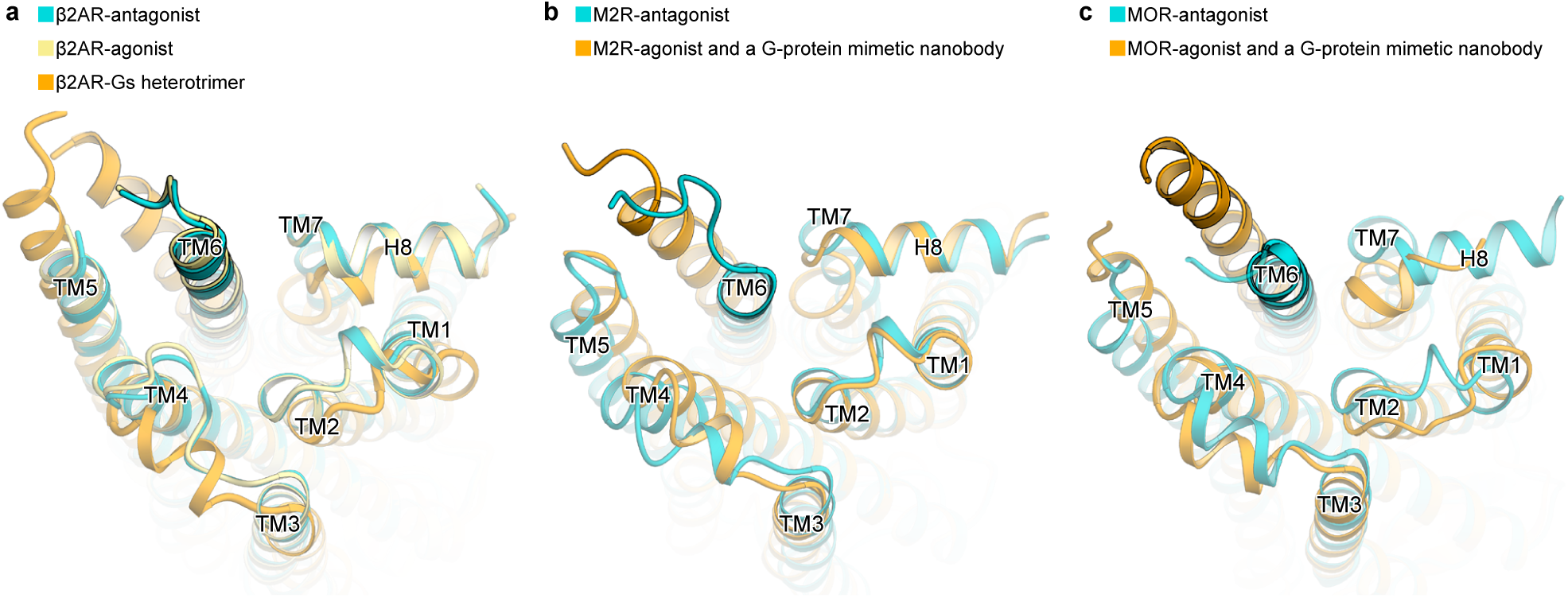
Comparison of the intracellular sides of class A GPCRs. **a**, Structures of β2 adrenergic receptor in complex with an antagonist (PDB 2RH1), an agonist (PDB 3PDS), and G-protein heterotrimer (PDB 3SN6), coloured turquoise, khaki, and orange, respectively. **b**, Structures of M2 receptor bound to an antagonist (PDB 3UON), and in complex with an agonist and G-protein mimetic nanobody (PDB 4MQS), coloured turquoise and orange, respectively. **c**, Structures of μ-opioid receptor bound to an antagonist (PDB 4DKL), and in complex with an agonist and a G-protein mimetic nanobody (PDB 5C1M), coloured turquoise and orange, respectively. Upon G-protein activation, the intracellular side of TM7 moves inward by 1 to 2 Å, and that of TM6 moves outward by 10 to 14 Å. These movements are a common structural feature among the class A GPCRs. Since agonist binding only induces structural heterogeneity on the intracellular side of TM6, the intracellular side of TM6 is not opened up in the agonist-bound structure of the β2 adrenergic receptor (PDB 3PDS). This process is highly dependent on the G-protein binding.

**Supplementary Fig. 6.**
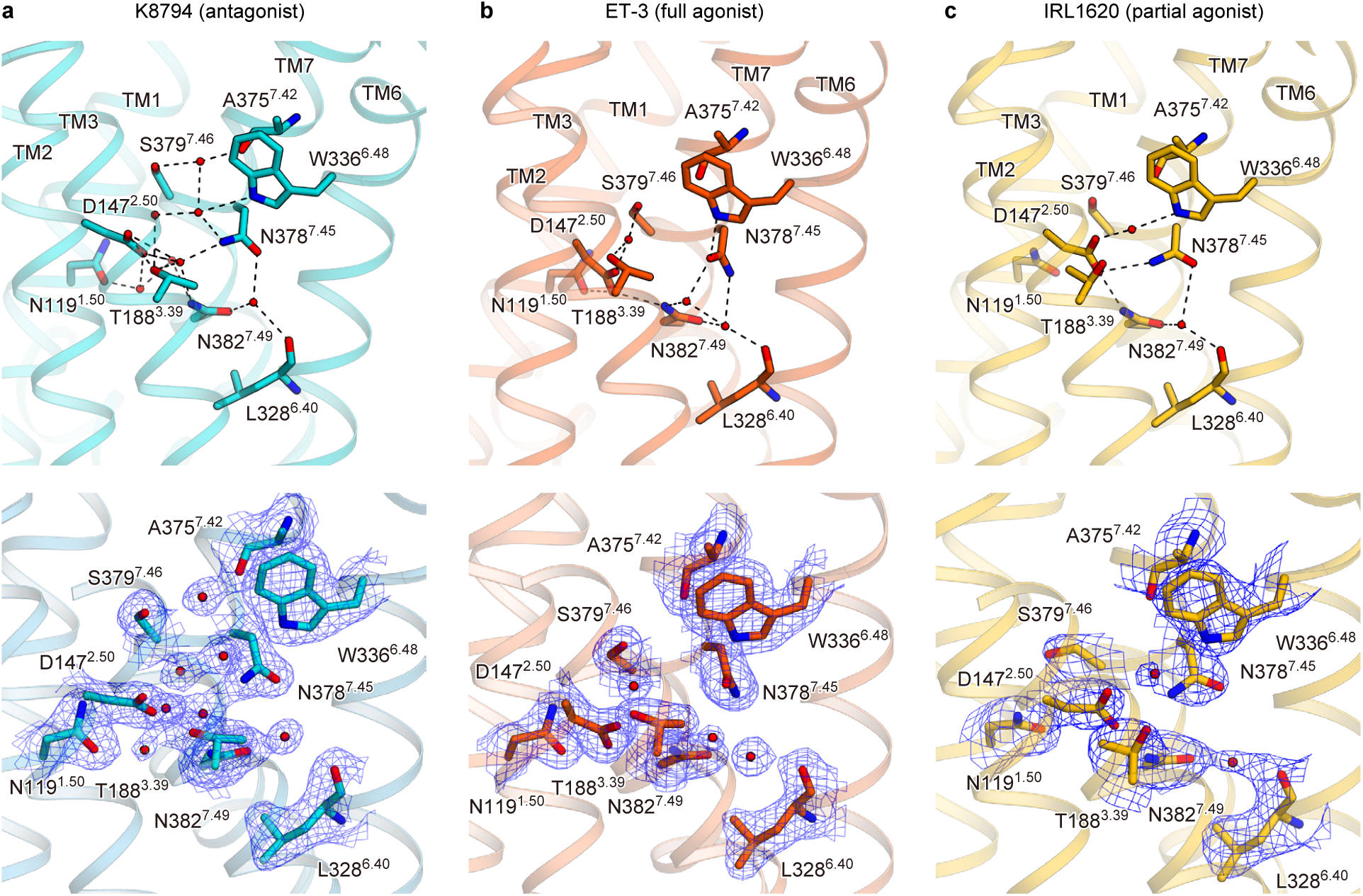
Water-mediated hydrogen-bonding network at the receptor core. **a-c**, Hydrogen-bonding networks in the intermembrane regions of the K8794- (**a**), ET-3-(**b**), and IRL1620- (**c**) bound structures, coloured as in Fig. 5. The upper panels show the overall hydrogen-bonding interactions. Waters are shown as red spheres, and hydrogen bonding interactions are indicated by dashed lines. The lower panels show the 2*F_o_* − *F_c_* maps around D147^2.50^ and W336^6.48^, contoured at 1.0σ.

**Supplementary Fig. 7.**
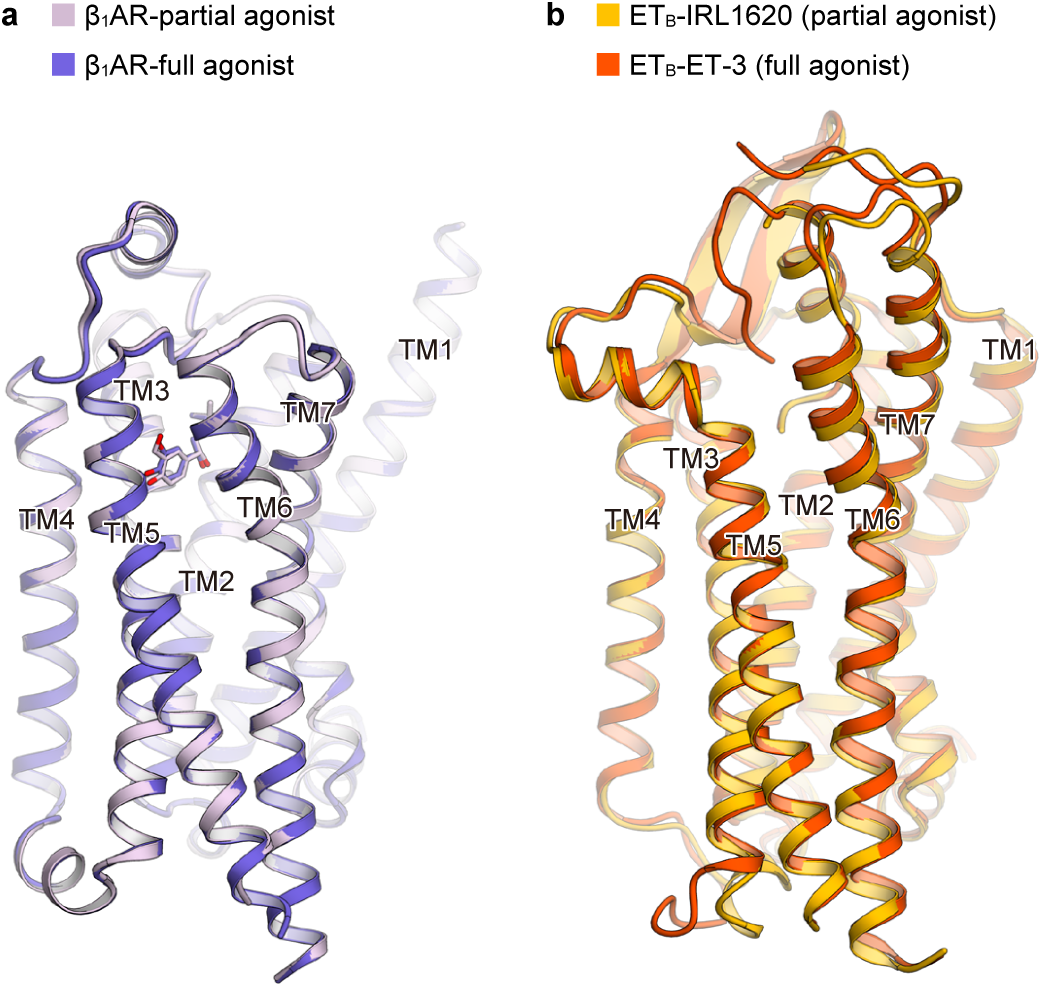
Comparison with β 1 adrenergic receptor structures. **a**, Superimposition of the β1 adrenergic receptor structures in complex with the full agonist isoprenaline (PDB 2Y03) and the partial agonist salbutamol (PDB 2Y04) (overall R.M.S.D of 0.25 Å for the C_α_ atoms). **b**, Superimposition of the ET_B_ structures in complex with the full agonist ET-3 and the partial agonist IRL1620 (overall R.M.S.D of 0.91 Å for the C_α_ atoms).

